# The Class I E3 Ubiquitin Ligase TRIM67 Modulates Brain Development and Behavior

**DOI:** 10.1101/241331

**Authors:** Nicholas P. Boyer, Caroline Monkiewicz, Sheryl S. Moy, Stephanie L. Gupton

## Abstract

Specific class I members of the TRIM family of E3 ubiquitin ligases have been implicated in neuronal development from invertebrates to mammals. The single invertebrate class I TRIM and mammalian TRIM9 regulate axon branching and guidance in response to the axon guidance cue netrin-1, whereas mammalian TRIM46 establishes the axon initial segment. In humans, mutations in TRIM1 and TRIM18 are implicated in Optiz Syndrome, characterized by midline defects and often mild intellectual disability. TRIM67 is the most evolutionarily conserved vertebrate class I TRIM, yet is the least studied. Here we show that TRIM67 interacts with both its closest paralog TRIM9 and the netrin receptor DCC, and is differentially enriched in specific brain regions at specific developmental points. We describe the anatomical and behavioral consequences of deletion of murine *Trim67*. While viable, mice lacking *Trim67* display severe impairments in spatial memory, cognitive flexibility, social novelty preference, muscle function and sensorimotor gating. Additionally, they exhibit abnormal anatomy of several brain regions, including the hippocampus, striatum and thalamus, as well as the corpus callosum. This study demonstrates the necessity for TRIM67 in appropriate brain development and function.

**SIGNIFICANCE STATEMENT:** As a family, class I TRIM E3 ubiquitin ligases play important roles in neuronal development and function, potentially cooperatively. TRIM67 is the most evolutionarily conserved class I TRIM and is developmentally regulated and brain-enriched. Deletion of murine *Trim67* results in malformations of a subset subcortical brain regions and of cortical and subcortical myelinated fiber tracts, as well as deficits in spatial memory, motor function, sociability and sensorimotor gating. We conclude that TRIM67 is critical for appropriate brain development and behavior, potentially downstream of the axon guidance cue netrin, and in cooperation with class I TRIM9.

## INTRODUCTION

E3 ubiquitin ligases mediate covalent attachment of ubiquitin to specific substrates. This posttranslational modification targets substrates for degradation or affects substrate localization, interactions, or function (Komander and Rape, 2012; Ciechanover, 2015). In developing neurons specific non-degradative ubiquitination events alter cytoskeletal dynamics and signaling pathways during axonal morphogenesis (Menon et al., 2015; Plooster et al., 2017). Although the expression patterns and functions of many ubiquitin ligases are not known, many are brain-enriched (Deshaies and Joazeiro, 2009) and implicated in neurodevelopmental and neurodegenerative diseases (Anuppalle et al., 2013; Atkin and Paulson, 2014). Specific members of the tripartite motif (TRIM) family of E3 ligases are brain-enriched and regulate neuronal development and function (Tocchini and Ciosk, 2015; Fatima et al., 2016; Olsson et al., 2016; Jin et al., 2017), however the functions and expression patterns of many TRIM proteins are unknown.

The TRIM family is subdivided into classes based on their carboxy-terminal domains (Short and Cox, 2006). The mouse and human genomes contain six class I TRIMs comprising three pairs of paralogs. Mutations in the TRIM1 and TRIM18 pair are associated with X-linked Opitz syndrome (Quaderi et al., 1997; Cainarca et al., 1999; Geetha et al., 2014), characterized by midline birth defects, and frequently intellectual and motor disabilities. TRIM46 is required for formation of axon initial segments (Van Beuningen et al., 2015), and mutations in its paralog TRIM36 are associated with anencephaly (Singh et al., 2017). TRIM9 and TRIM67 are the most evolutionarily conserved class I TRIMs (Short and Cox, 2006). TRIM9 localizes to Parkinsonian Lewy bodies (Tanji et al., 2010), is linked to atypical psychosis (Kanazawa et al., 2013), and regulates netrin-dependent axon guidance through the receptor DCC (Hao et al., 2010; Menon et al., 2015). *Trim9* deletion is also associated with aberrant migration, morphogenesis, and synapse organization of adult-born neurons in the murine dentate gyrus, and severe deficits in spatial learning and memory (Winkle et al., 2016). A single study has shown that TRIM67 is expressed in the mouse and human brain and regulates neuritogenesis in a mouse neuroblastoma cell line (Yaguchi et al., 2012), although its role in neurons and *in vivo* is unknown.

In *D. melanogaster* and *C. elegans* there is a single class I *Trim* (Short and Cox, 2006). Loss of this TRIM in either organism phenocopies the axon guidance and branching defects that occur upon loss-of-function of invertebrate orthologs of the axon guidance cue netrin or its receptor DCC (Alexander et al., 2010; Hao et al., 2010; Morikawa et al., 2011). Sequence comparison of the *Drosophila* and human class I TRIMs suggested that TRIM9 was the closest mammalian ortholog (Short and Cox, 2006; Morikawa et al., 2011). In mice, loss of *Ntn1* or *Dcc* causes agenesis of the corpus callosum and hippocampal commissure, as well as other axon guidance and branching defects (Serafini et al., 1996; Fazeli et al., 1997; Bin et al., 2015). In contrast to invertebrates, we have shown that deletion of *Trim9* does not phenocopy loss of *Ntn1* or *Dcc* and in some cases exhibits an opposite, gain-of-function phenotype, including increased axon branching and thickening of the corpus callosum (Winkle et al., 2014; Menon et al., 2015). This suggested TRIM9 was not the functional ortholog of the invertebrate class I TRIM.

Here we show that TRIM67 interacts with its paralog TRIM9 and the netrin receptor DCC. We found that deletion of murine *Trim67* results in impairments in spatial memory, cognitive flexibility, social novelty preference, muscle function and sensorimotor gating. Histological analysis demonstrated decreases in the size of several brain regions in *Trim67^-/-^* mice, including the hippocampus, caudate putamen, thalamus, and basolateral amygdala. Additionally, there were decreases in the size of the internal capsule and the dorsal commissures, including the corpus callosum and hippocampal commissure. The consequences of *Trim67* deletion on brain anatomy and behavior indicate that TRIM67 plays critical roles in brain development and function, which may occur downstream of netrin and DCC.

## METHODS

### Animals

Mouse lines were on a C57BL/7 background and were housed and bred at the University of North Carolina with approval from the Institutional Animal Care and Use Committee. Timed pregnant females were obtained by placing male and female mice together overnight; the following day was designated as E0.5 if the female had a vaginal plug. *Trim67^-/-^* mice were generated as previously described for the generation of the *Trim9^-/-^* line (Winkle et al., 2014). After Cre-mediated excision of exon 1 of *Trim67*, the subsequent 16 ATGs encode out-of-frame transcripts. This excision was confirmed by PCR genotyping.

### Reagents

Antibodies: TRIM67 rabbit polyclonal generated using murine TRIM67 recombinant protein aa45- 73 (used at 1:1000 for immunoblot, 1:500 for immunohistochemistry); TRIM9 rabbit polyclonal antibody (used at 1:1000 for immunoblot) (Winkle et al., 2014); mouse monoclonal against GAPDH (sc-166545, Santa Cruz Biotech (SCBT), used at 1:2500); mouse monoclonal against myc tag (sc-40, SCBT, used at 1:1000); mouse monoclonal against HA tag (05–904, Millipore, used at 1:1000); mouse monoclonal against β-III-tubulin (801202, BioLegend, used at 1:2000 for immunoblot, 1:1000 for immunohistochemistry). Fluorescent secondary antibodies labeled with Alexa Fluor 568 or Alexa Fluor 647 were obtained from Invitrogen.

The plasmid encoding myc-tagged human TRIM9 aa139–781 (myc-TRIM9ΔRING) was described previously (Winkle et al., 2014). The plasmid encoding myc-tagged mouse TRIM67 aa158–783 (myc-TRIM67ΔRING) was generated by cloning the partial mouse TRIM67 sequence into the pcs2 vector. pcDNA3-DCC-HA (HA-DCC) was acquired from Dr. Mark Tessier Lavigne (Rockefeller).

### Cell Culture and Western Blotting

E15.5 dissociated cortical neuron cultures were prepared as previously described (Viesselmann et al., 2011), and lysed at 2 days in vitro in modified RIPA buffer (50mM Tris-HCl, pH 7.5, 200mM NaCl, 0.5% NP-40, 2mM MgCh, 300mM sucrose, 15mM sodium pyrophosphate, 50mM NaF, 40mM β-glycerophosphate, 1mM sodium vanadate, 150μg/mL phenylmethanesulfonyl fluoride, 2mM dithiothreitol, 5mM N-ethylmaleimide, 3mM iodoacetamide, 2μg/mL leupeptin, 5μg/mL aprotinin). Protein from embryonic and adult tissues was obtained by dissection at E15.5 and P78, respectively, followed by lysis in modified RIPA buffer. SDS-PAGE and immunoblot analysis were performed using standard procedures with far-red-conjugated secondary antibodies (LI-COR Biosciences) imaged with an Odyssey Imager (LI-COR Biosciences).

### Immunoprecipitation

Coimmunoprecipitation assays were conducted according to standard procedures. Briefly: HEK293t cells were transfected using Lipofectamine 2000 (Invitrogen) by manufacturer protocol. Protein A/G beads (SCBT) coupled with a mouse anti-myc antibody were incubated with lysates overnight at 4°C. Beads were washed three times with lysis buffer and bound proteins were resolved by standard SDS-PAGE and immunoblot analysis.

### Elevated Plus Maze

Mice were given one five-minute trial on the plus maze, which had two walled arms (closed arms, 20cm in height) and two open arms. The maze was elevated 50cm from the floor, and the arms were 30cm long. Animals were placed on the center section (8cm × 8cm) and allowed to freely explore the maze. Measures were taken of time in, and number of entries into, the open and closed arms.

### Marble-bury Test

Mice were tested in a Plexiglas cage located in a sound-attenuating chamber with ceiling light and fan. The cage contained 5cm of corncob bedding, with 20 black glass marbles (14mm diameter) arranged in an equidistant 5X4 grid on top of the bedding. Subjects were given access to the marbles for 30 min. Measures were taken of the number of buried marbles (two thirds of the marble covered by the bedding).

### Olfactory Test

Several days before the olfactory test, an unfamiliar food (Froot Loops, Kellog Co.) was placed overnight in the home cages of the mice. Observations of consumption were taken to ensure that the novel food was palatable. Sixteen to twenty hours before the test, all food was removed from the home cage. On the day of the test, each mouse was placed in a large, clean tub cage (46cm L × 23.5cm W × 20cm H), containing paper chip bedding (3cm deep), and allowed to explore for five minutes. The animals was removed from the cage, and one Froot Loop was buried in the cage bedding. The animal was then returned to the cage and given fifteen minutes to locate the buried food. Measures were taken of latency to find the food reward.

### Hotplate Test

Individual mice were placed in a tall plastic cylinder located on a hotplate, with a surface heated to 55°C (IITC Life Science, Inc.). Reactions to the heated surface, including hindpaw lick, vocalization, or jumping, led to immediate removal from the hotplate. Measures were taken of latency to respond, with a maximum test length of 30sec.

### Open Field Assay

Mice were given a one-hour trial in an open field chamber (41cm × 41cm × 30cm) crossed by a grid of photobeams (VersaMax system, AccuScan Instruments). Counts were taken of the number of photobeams broken during the trial in five-minute intervals, with separate measures for locomotion (total distance traveled) and re△RING movements. Time spent in the center region of the open field was measured as an index of anxiety-like behavior.

### Acoustic Startle and Prepulse Inhibition

Subjects were given 2 acoustic startle tests (San Diego Instruments SR_Lab system), one at age 10–11 weeks, and another at age 17–19 weeks. Mice were placed in a small Plexiglas cylinder within a larger, sound-attenuating chamber. The cylinder was seated upon a piezoelectric transducer, which allowed vibrations to be quantified and displayed on a computer. The chamber included a ceiling light, fan, and a loudspeaker for the acoustic stimuli. Background sound levels (70dB) and calibration of the acoustic stimuli were confirmed with a digital sound level meter (San Diego Instruments).

Each session consisted of 42 trials that began with a five-minute habituation period. There were 7 different types of trials: the no-stimulus trials, trials with the acoustic startle stimulus (40msec; 120dB) alone, and trials in which a prepulse stimulus (20msec; either 74, 78, 82, 87 or 90dB) occurred 100ms before the onset of the startle stimulus. Measures were taken of the startle amplitude for each trial across a 65-msec sampling window, and an overall analysis was performed for each subject’s data for levels of prepulse inhibition at each prepulse sound level (calculated as: 100 - [(startle response with prepulse / startle response with no prepulse) × 100]).

### Accelerating Rotarod

In the first test session, mice were given three trials on an accelerating rotarod (Ugo Basile, Stoelting Co.) with 45 seconds between each trial. Two additional trials were given 48 hours later. Revolutions per minute was set at an initial value of 3, with a progressive increase to a maximum of 30 across 5 minutes (the maximum trial length). Measures were taken for latency to fall from the top of the rotating barrel.

### Gait Analysis

A track of Whatman filter paper (Sigma-Aldrich) was placed in a 7.5cm-wide, 40cm-long Plexiglas corridor closed at one end and open to a 10×10 cm Plexiglas chamber at the other end. The chamber was covered with an opaque cloth and filled with several pellets of dry food. Mouse forepaws and hindpaws were painted with two colors of Crayola Washable Kids’ Paint (Crayola, LLC) before the mouse was placed at the closed end of the Plexiglas corridor. Once mice had walked the length of the corridor, they were given 45 seconds in the recovery chamber before being returned to their home cage.

### Rolling Wire-hang

Mice were held near a circular rubber gasket, suspended from a pulley, such that the mouse grasped the loop (Hoffman and Winder, 2016). Mice were released once all four paws were engaged, and latency to fall was recorded in three trials separated by 45 seconds each. Latency to fall was measured in seconds, and then multiplied by weight to produce hang impulse.

### Three-Chamber Sociability Assay

This procedure consisted of 3 10-minute phases: a habituation period, a test for sociability, and a test for social novelty preference. For the sociability assay, mice were given a choice between being in the proximity of an unfamiliar, sex-matched C57BL/6J adult mouse (“stranger 1”) versus being alone. In the social novelty phase, mice were given a choice between the already-investigated stranger 1, versus a new unfamiliar mouse (“stranger 2”). The social testing apparatus was a rectangular, 3-chambered box fabricated from clear Plexiglas. Dividing walls had doorways allowing access into each chamber. An automated image tracking system (Noldus Ethovision) provided measures of time spent within 5cm of the Plexiglas cages (the cage proximity zone), and entries into each side of the social test box.

At the start of the test, the mouse was placed in the middle chamber and allowed to explore for 10 minutes, with the doorways into the 2 side chambers open. After the habituation period, the test mouse was enclosed in the center compartment of the social test box, and stranger 1 was placed in one of the side chambers. The stranger mouse was enclosed in a small Plexiglas cage drilled with holes, which allowed nose contact. An identical empty Plexiglas cage was placed in the opposite side chamber. Following the placement of the stranger and the empty cage, the doors were re-opened, and the subject was allowed to explore the social test box for a 10-minute session. At the end of the sociability phase, stranger 2 was placed in the empty Plexiglas container, and the test mouse was given an additional 10 minutes to explore the social test box.

### Morris Water Maze

The water maze consisted of a large circular pool (diameter = 122cm) partially filled with water (45cm deep, 24–26°C), located in a room with numerous visual cues. The procedure involved three different phases: a visible platform test, acquisition in the hidden platform task, and reversal learning. In the visible platform test, each mouse was given 4 trials per day, across 2 days, to swim to an escape platform cued by a patterned cylinder extending above the surface of the water. For each trial, the mouse was placed in the pool at 1 of 4 possible locations (randomly ordered), and then given 60 seconds to find the visible platform. If the mouse found the platform, the trial ended, and the animal was allowed to remain 10 seconds on the platform before the next trial began. If the platform was not found, the mouse was placed on the platform for 10 seconds, and then given the next trial. Measures were taken of latency to find the platform and swimming speed via an automated tracking system (Noldus Ethovision).

For the hidden platform task, the platform (diameter = 12cm) was submerged. Each animal was given 4 trials per day, with 1 minute per trial, to swim to the hidden platform. Criterion for learning was an average group latency of 15 seconds or less to locate the platform. Mice were tested until the group reached criterion, with a maximum of 9 days of testing. When criterion was reached, mice were given a one-minute probe trial in the pool with the platform removed. Selective quadrant search was evaluated by measuring percent time spent in the target quadrant and number of crossings over the location where the platform (the target) had been placed during training, versus the corresponding area in the opposite quadrant. During the subsequent reversal learning phase, the escape platform was placed in a different quadrant, and mice were tested until the group reached the learning criterion, using the same procedure as with initial acquisition. A second one-minute probe trial was conducted at the end of the reversal learning phase.

### Brain Fixation and Sectioning

All adult mice used for neuroanatomical studies were anesthetized with an intraperitoneal injection of 1.2% avertin and intracardially perfused with 1X PBS followed by 4% paraformaldehyde (PFA). Brains were removed and fixed in 4% PFA for an additional 48 hours, rinsed with 1X PBS and washed in 70% EtOH for at least 24 hours prior to vibratome sectioning at 100 μm thickness. Embryonic brains were removed at E15.5 and drop-fixed in 4% PFA for 72 hours, rinsed with 1X PBS, and embedded in 1% agarose prior to cryoprotection with 30% sucrose and cryostat sectioning at 20μm thickness.

### Immunohistochemistry

Sections were blocked in 0.2% Triton-X 100 and 10% BSA in 1X PBS for 2 hours on a rocker at room temperature. Solution was replaced with primary antibodies in blocking solution overnight at 4°C. Sections were washed three times for 20 minutes each in 1X PBS, and then incubated with secondary antibodies in blocking solution for 2 hours at room temperature. Sections were then incubated with 300nM DAPI in 1X PBS for 30 minutes, and then washed three times for 20 minutes each with 1X PBS. Sections were mounted in mounting media (20mM Tris pH 8.0, 90% glycerol, 0.5% N-propyl gallate) and sealed with nail polish. Fluorescently labeled sections were imaged on a Zeiss LSM 780 confocal microscope using a 20X Plan-Apochromat objective.

### Histological Measurements

Five-week-old brains from an equal number of male and female mice from both genotypes were sectioned and stained with black gold II (Histo-Chem, Inc.) per manufacturer protocol and mounted in DPX mounting media (VWR International), and imaged on a Leica WILD M420 macroscope using an APOZOOM lens. Sections were aligned to a reference atlas, and individual regions were outlined and measured using ImageJ. White matter in the striatum was defined using the Trainable Weka Segmentation v3.2.2 plugin for ImageJ (Arganda-Carreras et al., 2017) trained on 2 classes (white and gray matter) with at least 8 examples per class. Brain regions and fiber tracts were measured in each section, then individually aligned by minimizing the total standard deviation, as the shapes of regions along the anterior-posterior axis were not apparently different between individual animals.

### Protein Sequence Comparison

Protein sequences for TRIM9 and TRIM67 were retrieved from the UniprotKB/Swiss-Prot (accession numbers: Q9C026–1, Q8C7M3–3, Q6NZX0, F7E188, A0A1D5Q2P9, Q91ZY8, A0A1D5PZ60, M9MRI4, B1GRL4, Q6ZTA4–3, Q505D9–1, A0A286YAS5, F6U581, F6S2G3, D3ZTX1, A0A1L8F061, A0A1L8G7R8) and NCBI GenPept (accession numbers: XP_015139864.1, EKC37550.1) databases. Sequences were compared using the Clustal Omega multiple sequence alignment, with default parameters (Sievers et al., 2011).

### Statistical Methods

Behavioral data were analyzed using one-way or repeated measures Analysis of Variance (ANOVA), with factor genotype. Separate ANOVAs for each sex were conducted for measures of body weight. Fisher’s protected least-significant difference (PLSD) tests were used for comparing group means only when a significant F value was determined Within-group comparisons were conducted to determine side preference in the three-chamber test for social approach, and for quadrant preference in the Morris water maze.

For all brain regions measured in multiple sections, comparisons were performed by two-way ANOVA. Sections and genotypes were treated as groups with each brain a subject, with genotype as the factor. For the paired brain weight comparison, a Wilcoxon signed-rank test was used. A chi-square test was used to compare the heterozygote birth rate to the predicted Mendelian distribution. For all comparisons, α = 0.05.

## RESULTS

### TRIM67 is evolutionarily conserved and interacts with DCC and TRIM9

Comparisons of human class I TRIMs to *D. melanogaster* trim9 suggested that human TRIM9 was the most evolutionarily conserved (Short and Cox, 2006; Morikawa et al., 2011). In light of the divergence in phenotypes associated with the loss-of-function mutations in invertebrate trim9 compared to deletion of murine *Trim9* (Hao et al., 2010; Morikawa et al., 2011; Winkle et al., 2014; Menon et al., 2015), we compared the sequences of invertebrate class I TRIMs to the class I TRIMs from several vertebrate genomes. Consistent with other analysis, TRIM67 and TRIM9 were the most conserved vertebrate class I TRIMs. In addition, this broader phylogenetic analysis suggested that vertebrate TRIM67 homologs were more evolutionarily conserved than vertebrate TRIM9 homologs **(Fig. 1*A***).

**Fig. 1:**
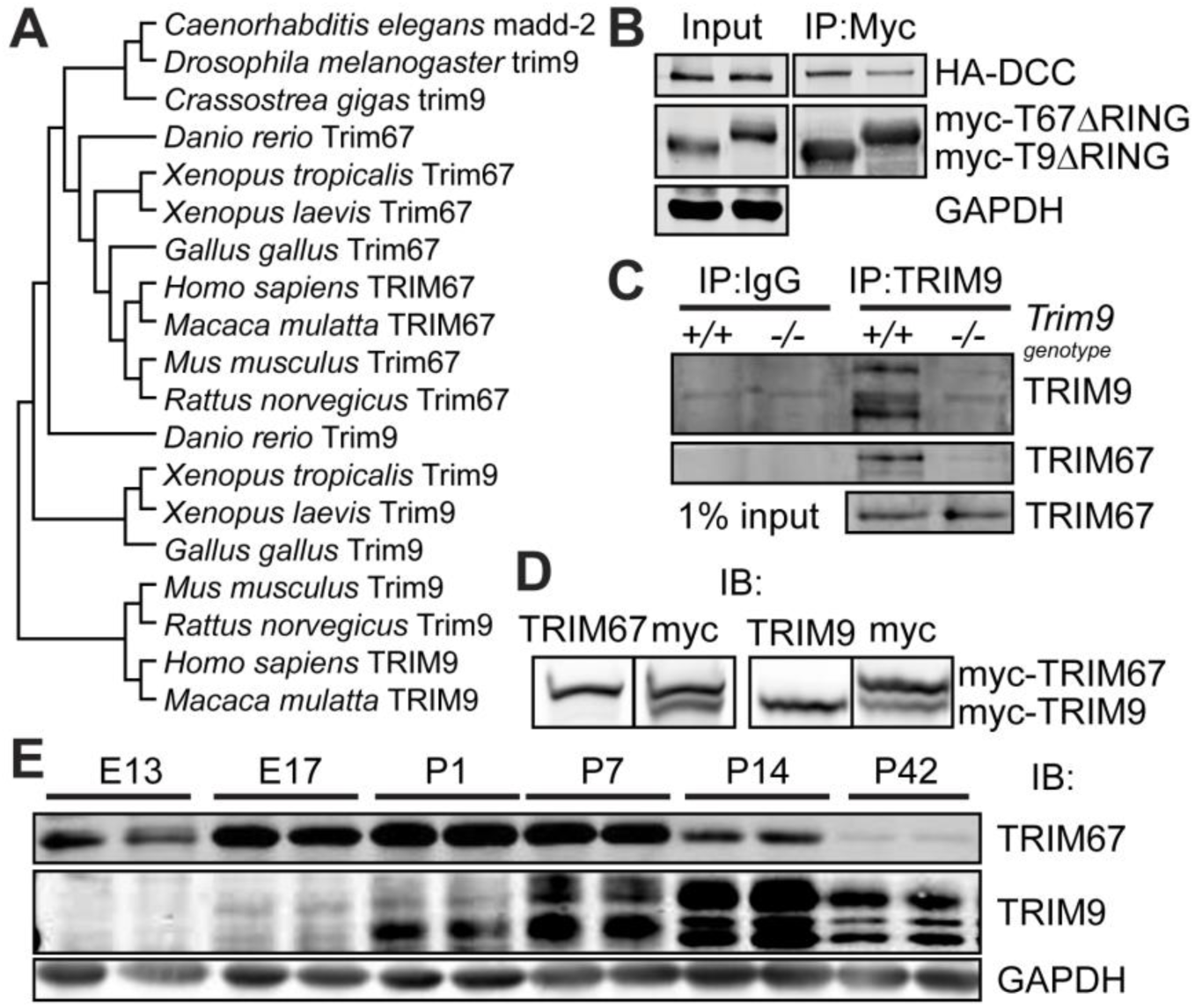
TRIM67 is evolutionarily conserved and interacts with DCC and TRIM9. **A** Phylogenetic tree of invertebrate class 1 TRIM proteins alongside several vertebrate TRIM9 and TRIM67 homologs. Vertebrate TRIM67s exhibit higher sequence similarity to invertebrate class 1 TRIMs. **B** HEK293t cells were co-transfected with HA-tagged DCC and myc-tagged TRIM9 or TRIM67 lacking the RING domain (△RING). Immunoprecipitation of either myc-TRIM△RING coprecipitated HA-DCC. **C** Western blot of TRIM9 immunoprecipitates from *Trim9^+/+^ and Trim9^-/-^* mouse cortical neuronal lysate probed for TRIM9 and TRIM67. Co-immunoprecipitation of TRIM67 occurs only in the presence of TRIM9. **D** Images of duplicate western blot lanes of lysate from HEK293 cells expressing both myc-TRIM9 and myc-TRIM67, and probed with both myc (right) and the indicated TRIM antibody (left). The newly generated TRIM67 polyclonal antibody recognizes TRIM67 but not TRIM9; the polyclonal TRIM9 antibody recognizes TRIM9, but not TRIM67. **E** Western blots of cortical lysate from indicated embryonic (E) and postnatal (P) ages reveal expression patterns of TRIM67 and the three isoforms of TRIM9.

Mammalian and invertebrate TRIM9 interact with the netrin-1 receptor DCC and regulate netrin-dependent axonal responses (Hao et al., 2010; Morikawa et al., 2011; Winkle et al., 2014). Co-immunoprecipitation assays with HA-tagged DCC and either myc-tagged TRIM67△RING or myc-TRIM9△RING from HEK293 lysates demonstrated that both TRIM9 and TRIM67 interact with DCC **(Fig. 1*B***). These constructs were used because deletion of the RING domain (△RING) stabilizes interactions between TRIMs and their binding partners (Kim et al., 2015). TRIM proteins often heterodimerize, including class I members TRIM1 and TRIM18 (Hatakeyama, 2011). We exploited co-immunoprecipitation assays to determine if TRIM9 and TRIM67 heterodimerized. Endogenous TRIM67 was co-immunoprecipitated by a polyclonal TRIM9 antibody from wildtype embryonic cortical lysate, but not from *Trim9^-/-^* lysate **(Fig. 1*C***), suggesting TRIM9 and TRIM67 heterodimerize. Using specific polyclonal antibodies that we developed against unique regions within the N-terminus of TRIM67 and TRIM9 **(Fig. 1*D***), we found both were expressed in the murine cortex at a range of ages. TRIM67 expression peaks embryonically and perinatally, whereas expression of TRIM9 peaks later in development **(Fig. 1*E***).

### Generation of *Trim67^-/-^* Mice

We previously generated mice in which the first exon of *Trim9*, which encodes the RING and two BBox domains, was flanked by LoxP sites (Winkle et al., 2014). Exon 1 was removed from the germline with cytomegalovirus (CMV)-Cre, and all TRIM9 protein was lost. Since *Trim9* and *Trim67* have similar gene architectures, we generated a conditional *Trim67* allele **(Fig. 2*A***, *Trim67^fl^)* by flanking exon one and ~200 bp upstream of the ATG start site by LoxP sites via homologous recombination. Mice carrying the *Trim67^fl^* allele were crossed with CMV-Cre mice to delete *Trim67* in the germline and generate *Trim67^-/-^* mice. Multiplexed PCR genotyping confirmed deletion of exon 1 of *Trim67* **(Fig. 2*B***). *Trim67^-/-^* mice are viable, and heterozygous adults produced the expected Mendelian ratio of offspring **(Fig. 2*C***). Western blotting of lysates from embryonic whole-brain and dissociated cortical neuron culture at two days *in vitro* confirmed loss of TRIM67 protein **(Fig. 2*D***).

**Fig. 2:**
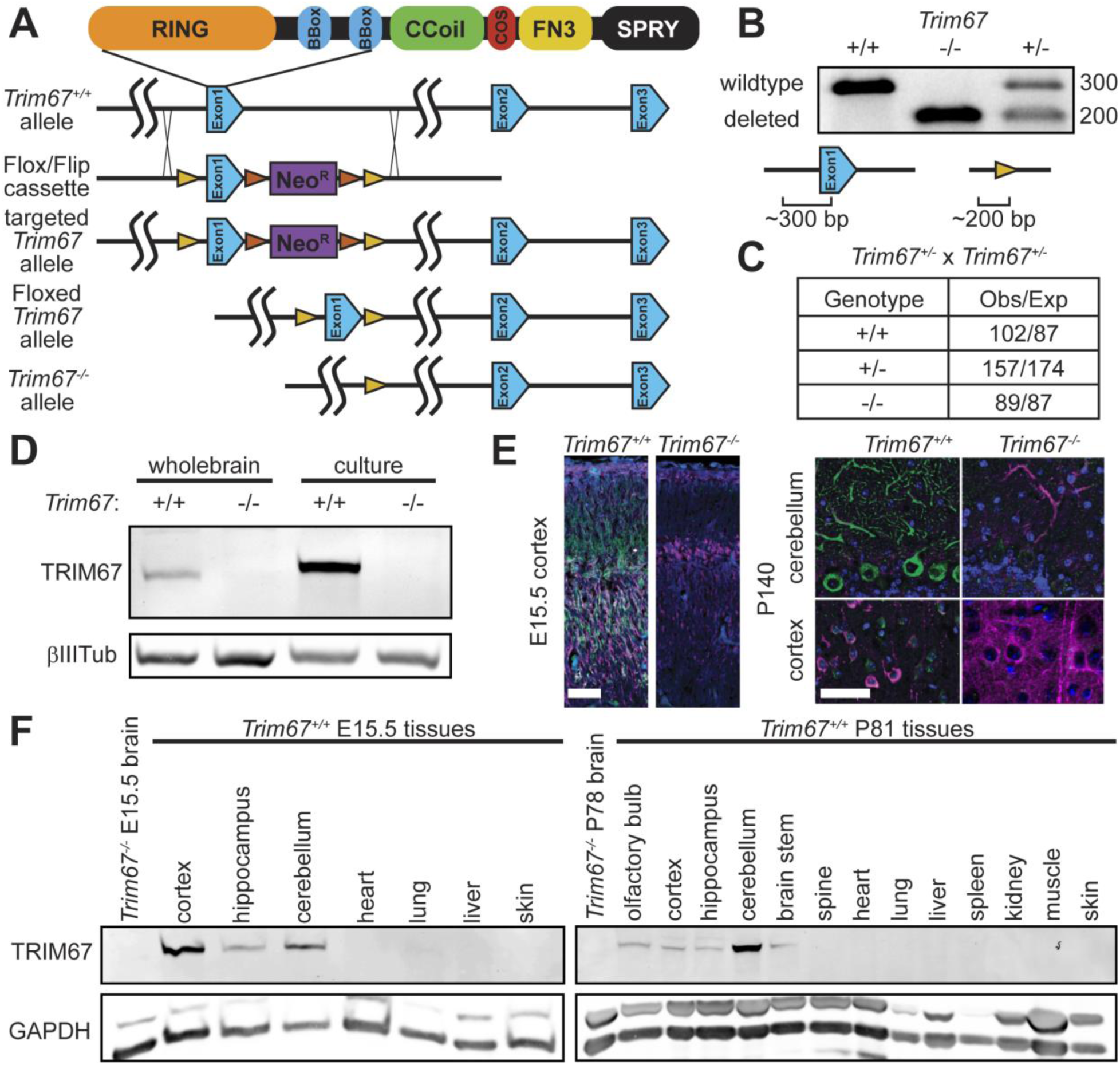
Generation of *Trim67^-/-^* mouse and TRIM67 brain localization. **A** Diagram of targeting strategy for knockout of the *Trim67* gene showing Cre-lox mediated excision of exon 1. This excision leads to the next 16 intronic and 4 exonic ATG codons being out of frame. LoxP sites are shown in yellow, FLIP sites in orange, and neomycin resistance gene (NeoR) in purple. **B** Agarose gel separation of genotyping PCR products, demonstrating deletion of both copies of the first exon of *Trim67* (*-/-*) or one copy in a heterozygote (-/+). Diagrams show PCR products from *Trim67^+/+^* and *Trim67^-/-^* DNA. **C** Mendelian ratio of *Trim67* allele inheritance in 48 litters of heterozygous knockout crosses, showing approximately expected rates of inheritance of each allele (p = 0.117, chi-square test). **D** Western blot of whole-brain or two day *in vitro* dissociated cortical neuron lysate from *Trim67^+/+^* and *Trim67^-/-^* E15.5 embryos probed for TRIM67 and βIII-tubulin as a loading control. **E** Fluorescent micrographs of E15.5 and P140 brain sections shows expression and loss of TRIM67 (green) in *Trim67^+/+^* and *Trim67^-/-^* mice, respectively. Sections are counterstained for β-III-tubulin (magenta) and with DAPI (blue). Scale bar 50μm. **F** Western blot detects TRIM67 expression in various adult and embryonic brain tissues, but not outside the nervous system. GAPDH is a loading control.

Immunohistochemical staining of embryonic brains demonstrated high TRIM67 expression throughout the developing cortex **(Fig. 2*E***), the hippocampus, and all structures of the nascent diencephalon **(Fig. 3*A,B,C***). This staining was absent in *Trim67^-/-^* embryonic brains. In the adult brain, TRIM67 was enriched in Purkinje cells of the cerebellum, and was evident in layers 2 and 3 of the cortex **(Fig. 2*E***), cortical axon tracts passing through the caudate putamen (CPu), and scattered cells within the midbrain **(Fig. 3*D,E***, arrowheads), but TRIM67 was not detected in the hippocampus **(Fig. 3*F***). Although TRIM67 was detected in most regions of the adult and embryonic brain, it was not detected in other tested tissues **(Fig. 2*F***) consistent with a previous report (Yaguchi et al., 2012). Compatible with immunohistochemistry, TRIM67 protein levels in the embryo were high in the cortex, whereas in the adult brain they were high in the cerebellum.

**Fig. 3:**
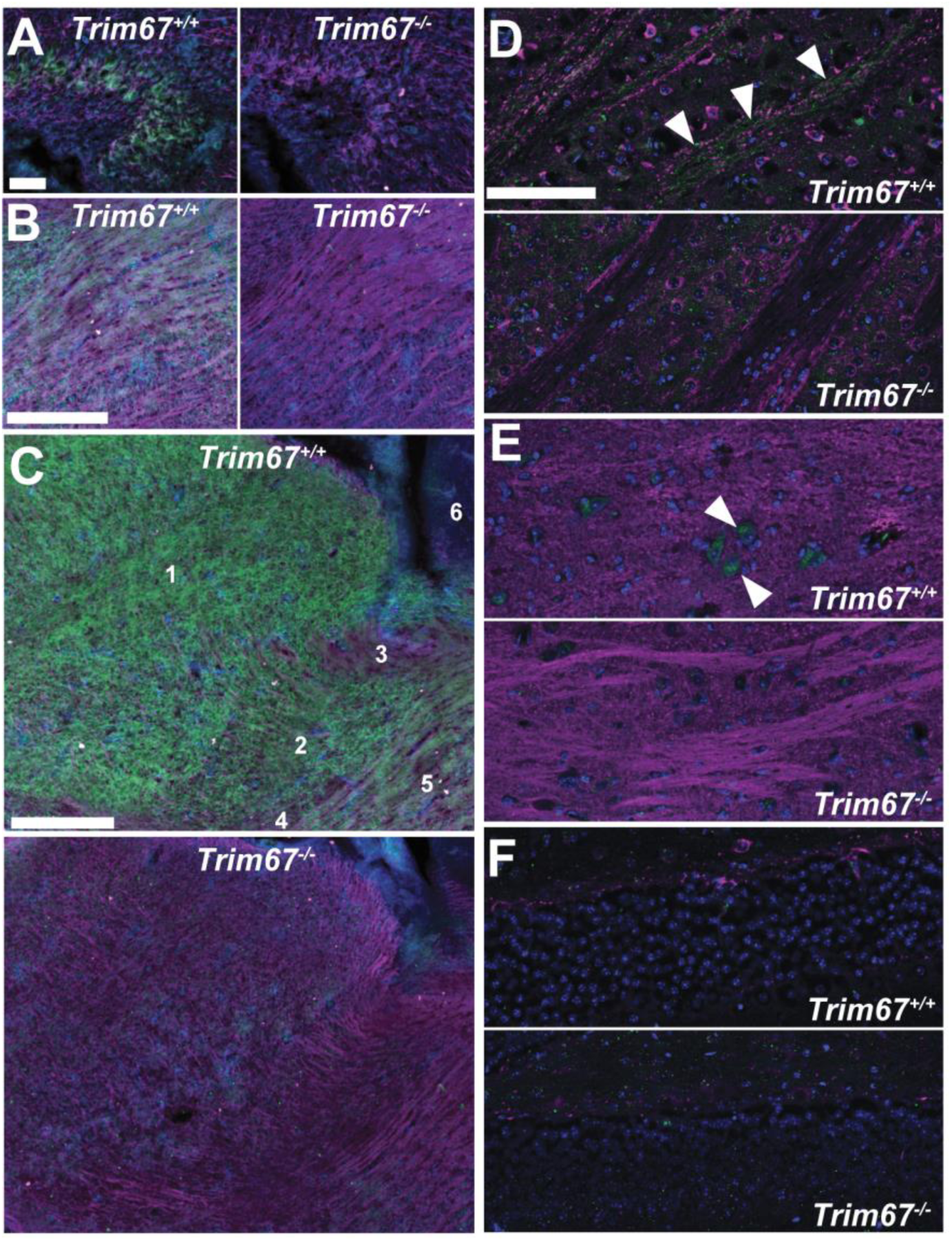
TRIM67 is expressed in multiple murine brain regions. Sagittal sections of E15.5 (**A-C**) and P140 (**D-F**) brains stained for TRIM67 (green), β-III-tubulin (red), and nuclei (DAPI, blue). **A** TRIM67 is expressed in cell bodies in the developing hippocampus at E15.5 (scale bar = 50 μm). **B,C** TRIM67 expression is evident in the peduncular hypothalamus (B, C5), diencephalon (C1), and reticular complex (C2), but not in the prethalamic eminence (C3), zona incerta complex (C4) or subpallium (C6) (scale bars B,C = 200μm). **D-F** In the adult brain, TRIM67 expression is evident in fiber bundles of the corticospinal tract passing through the caudate putamen (arrowheads, D) and scattered cells in the lateral hypothalamic area (arrowheads, **E**), but not in the hippocampus (**F**). (scale bar = 100μm).

### Cortical Fiber Tracts

The conservation of TRIM67, its embryonic expression in the cortex and hippocampus, and its interaction with DCC **(Fig1** *C*), along with the involvement of the invertebrate class 1 TRIM in axon guidance in the netrin-1/DCC pathway suggested that TRIM67 may function in the formation of netrin-dependent fiber tracts. We measured the size of several axon tracts in adult *Trim67^+/+^* and *Trim67^-/-^* littermate pairs **(Fig. 4*A***). Loss of murine *Ntn1* or *Dcc* leads to a narrower anterior commissure and agenesis of the corpus callosum and hippocampal commissure (Serafini et al., 1996; Fazeli et al., 1997). Deletion of *Trim67* had no effect on the size of the anterior commissure **(Fig. 4*B***) or the anterior corpus callosum, but did decrease the thickness of the posterior portion of the corpus callosum located above the hippocampus **(Fig. 4*C***). Additionally, the hippocampal commissure, measured as the combined dorsal fornix and dorsal hippocampal commissure, was thinner in *Trim67^-/-^* mice **(Fig. 4*C***). Taken together, these data suggest loss of *Trim67* decreases the size of cortical and hippocampal axon tracts, but not all commissures.

**Fig. 4:**
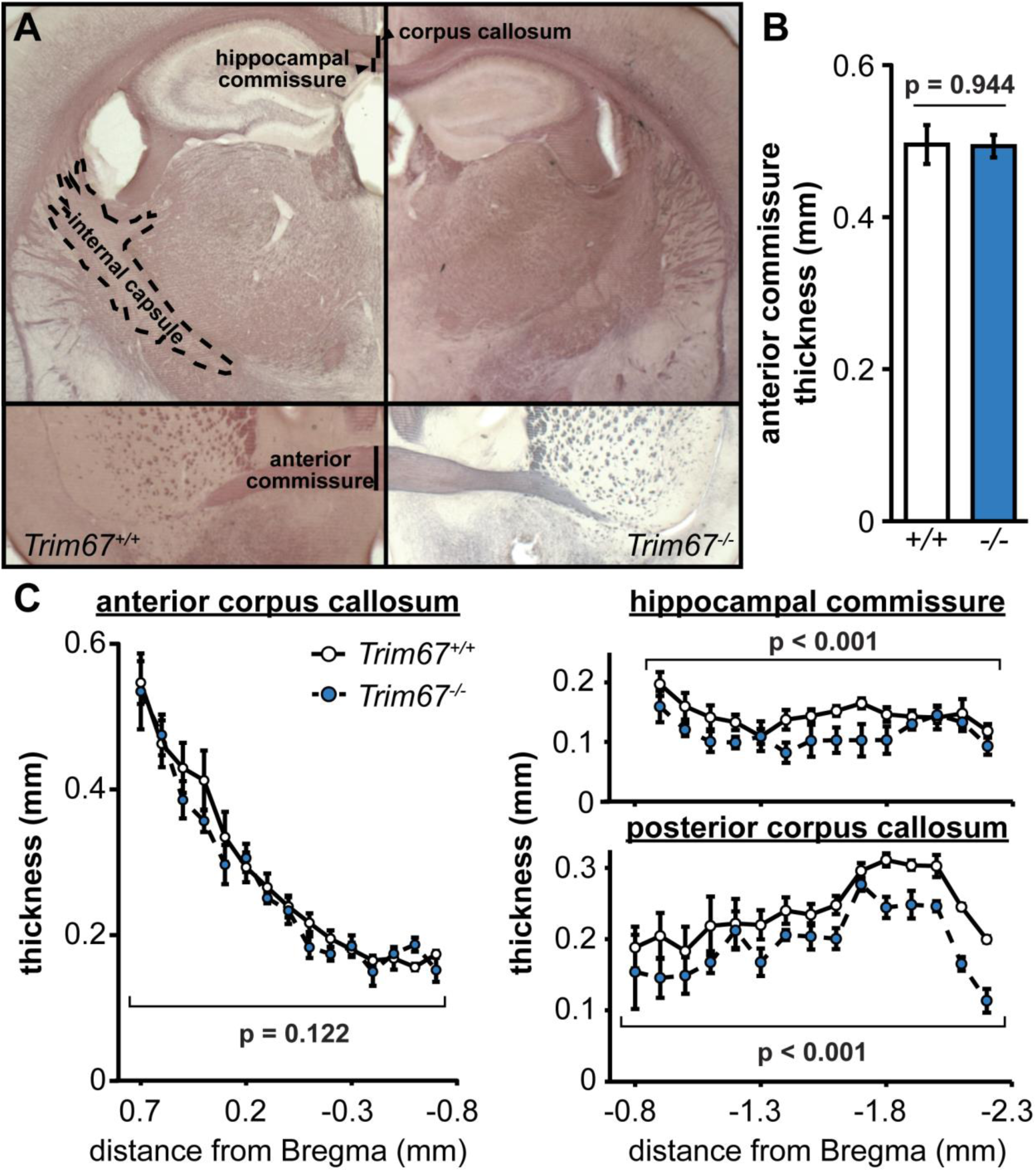
*Trim67* deletion affects the thickness of certain cortical fiber tracts. **A** Photomicrographs of black-gold stained coronal brain sections indicating fiber tracts used for measures. **B** Width of the thickest region of the anterior commissure was not affected by *Trim67* deletion (p = 0.944). **C** The thickness of the anterior corpus callosum was unaffected by *Trim67* deletion (p = 0.1218), however the posterior corpus callosum was thinner in *Trim67^-/-^* mice (p = 0.000110). The hippocampal commissure, including both the fornix and dorsal hippocampal commissure, was reduced in thickness in *Trim67^-/-^* mice (p = .000215).

### Behavioral Phenotyping of *Trim67^-/-^* Mice

In light of these fiber tract defects, we employed a battery of behavioral assays to determine the ramifications of loss of *Trim67 in vivo*. The regimen of behavioral testing (Table 1) has been standardized across multiple mouse strains (Moy et al., 2007, 2008, 2012; Huang et al., 2013; Nagy et al., 2017). Over the course of the behavioral testing, there was no significant effect of loss of *Trim67* on body weight or overall growth in either male or female mice **(Fig. 5*A***). There were also no effects of genotype on anxiety-like behavior in an elevated plus maze, exploratory and perseverative digging in a marble-bury assay, olfactory ability in finding buried food, or thermal sensitivity **(Fig. 5*B***). In an open field assay, loss of *Trim67* had no effect on locomotor activity or rearing movements **(Fig. 5*C***) or in time spent in the center region of the field *(Trim67^+/+^* = 234 ± 47s, *Trim67^-/-^* = 213 ± 32s), a measure of anxiety.

**Table 1:** Timecourse of behavioral assays conducted on *Trim67^+/+^* and *Trim67^-/-^* littermates.

| Age (weeks) | Procedure |
| --- | --- |
| 6-7 | Elevated plus maze test for anxiety-like behavior |
| 7-8 | Activity in a 1-hour open field test |
| 8-9 | Rotarod test for motor coordination and motor learning |
| 9-10 | Social approach in a three-chamber choice task |
| 10-11 | Marble-bury assay for digging responses; prepulse inhibition of acoustic startle responses (first test) |
| 11-12 | Buried food test for olfactory ability |
| 12-17 | Morris water maze test for spatial learning |
| 17-19 | Prepulse inhibition of acoustic startle responses (second test); hot-plate test for thermal sensitivity |

**Table 2:**
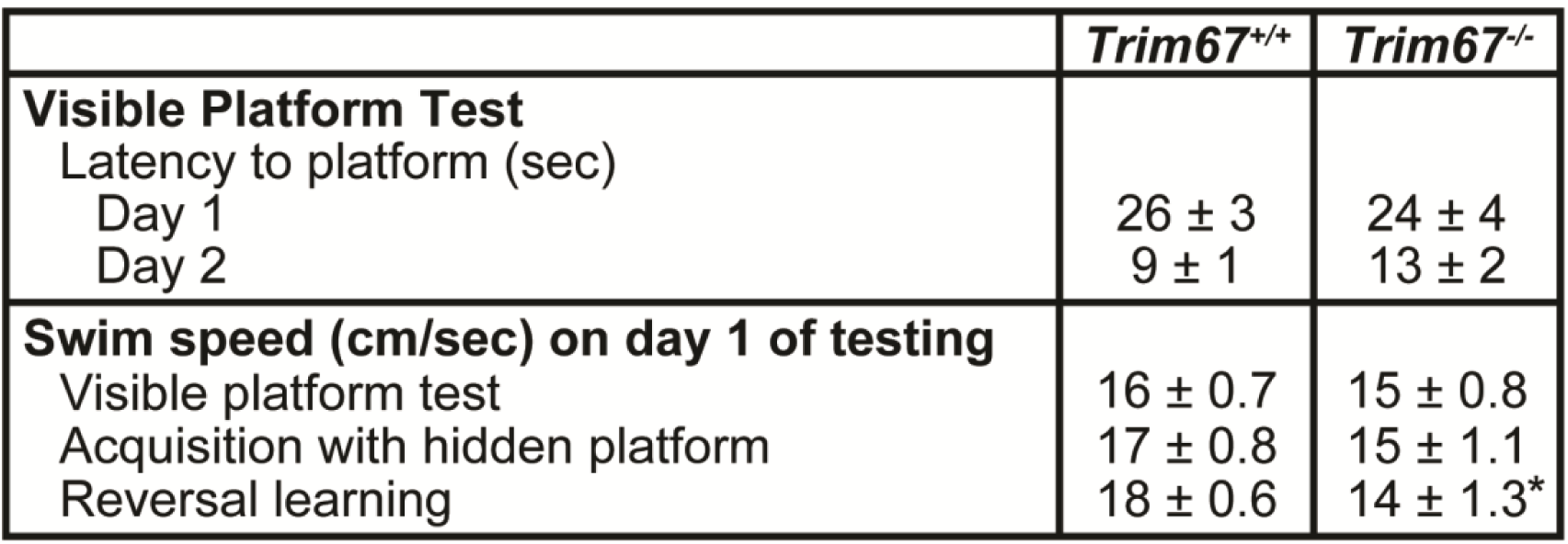
Swim speeds of mice during Morris water maze trials.

|  | *Trim67<sup>+/+</sup>* | *Trim67<sup>-/-</sup>* |
| --- | --- | --- |
| **Visible Platform Test** |  |  |
| Latency to platform (sec) |  |  |
| Day 1 | 26 ± 3 | 24 ± 4 |
| Day 2 | 9 ± 1 | 13 ± 2 |
| **Swim speed (cm/sec) on day 1 of testing** |  |  |
| Visible platform test | 16 ± 0.7 | 15 ± 0.8 |
| Acquisition with hidden platform | 17 ± 0.8 | 15 ± 1.1 |
| Reversal learning | 18 ± 0.6 | 14 ± 1.3* |

**Fig. 5:**
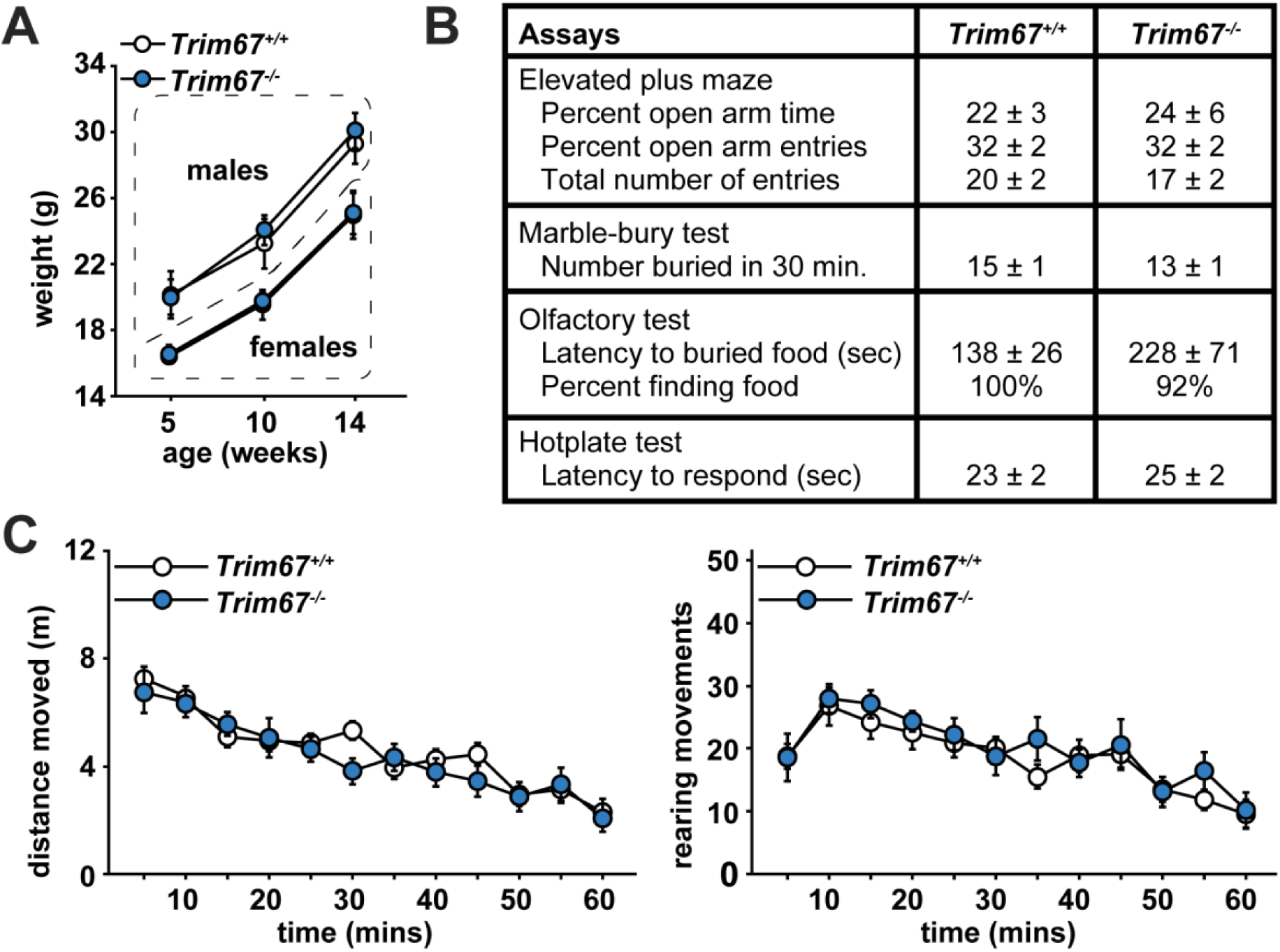
Mouse growth, sensory ability and general locomotion are not affected by *Trim67* deletion. **A** Weights of mice over the course of behavioral assays, showing no effect of *Trim67* loss on overall growth. **B** Results of elevated plus maze and several sensory assays. Units are indicated. **C** Locomotor results in the open field assay. None of these assays show significant results by ANOVA.

### *Trim67^-/-^* Mice Have Impaired Muscle Function

Mice carrying a spontaneous mutation in *Dcc* display overt impairments in motor function, including hopping behavior instead of a typical gait pattern (Finger et al., 2002; Welniarz et al., 2017). Assessment of footprints on a linear track **(Fig. 6*A***) indicated there were no differences between stride length or in either front or rear base width for *Trim67^+/+^* and *Trim67^-/-^* mice, suggesting no effect on gait **(Fig. 6*B***). Subjects were tested for motor coordination and learning on an accelerating rotarod. In the initial day of testing (trials 1–3) *Trim67^-/-^* mice exhibited deficits in acquisition of motor learning **(Fig. 6*C***), as indicated by a decreased latency to fall from the top of the rotating barrel. This difference was not observed on the second day of testing 48 hours later (trials 4,5). This suggests that deletion of *Trim67* leads to impairment of the initial performance, but not eventual learning, of a motor task. Since motor learning was not impaired, we suspected that decreased initial latency to fall may be due to impaired muscle function, and assayed muscle strength in a four-paw rolling-wire hang (Hoffman and Winder, 2016). The maximum and average hang impulse of three trials were significantly lower in *Trim67^-/-^* mice **(Fig. 6*D***), suggesting an overall decrease in muscle tone.

**Fig. 6:**
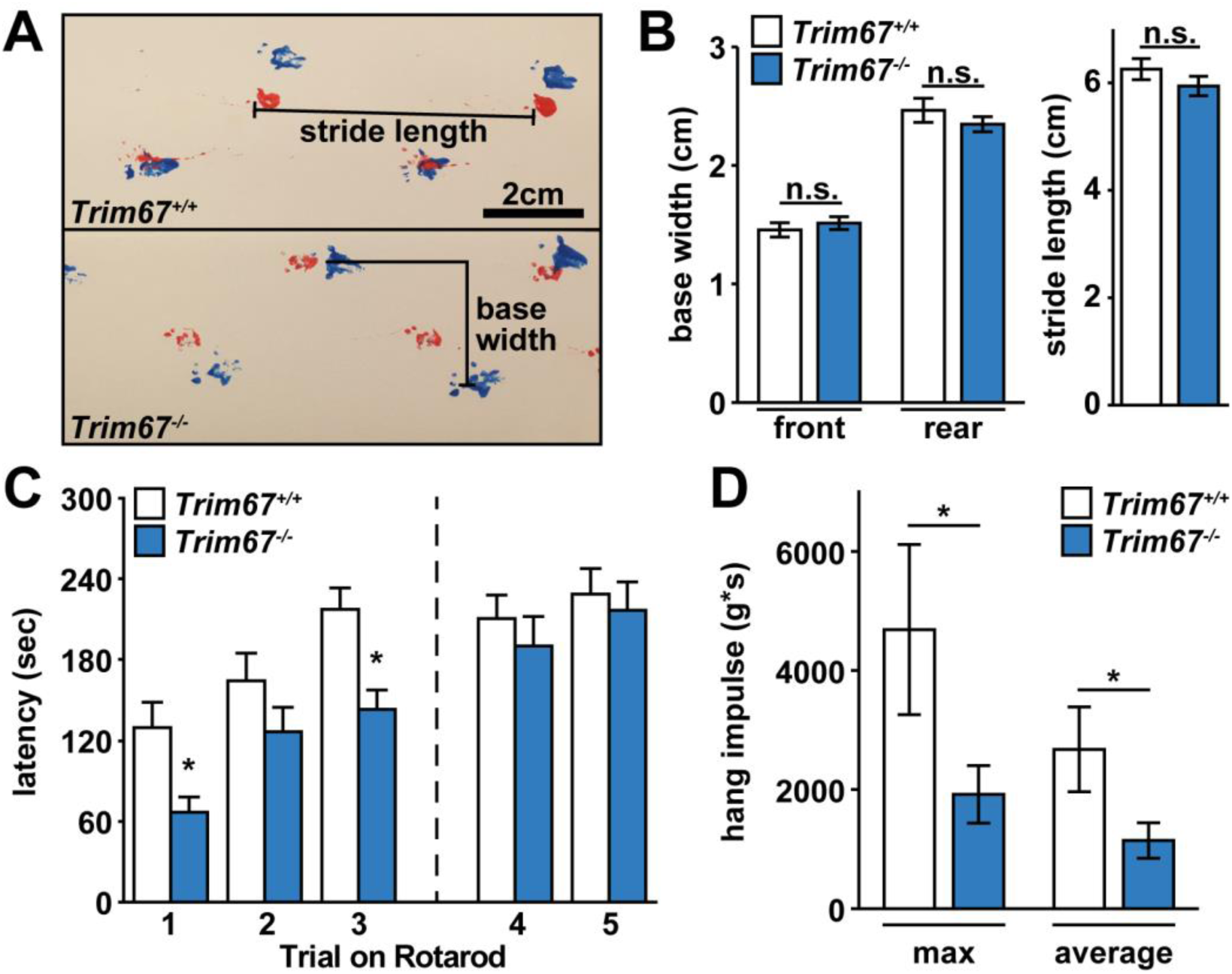
Muscle function, but not motor learning, is impaired in *Trim67^-/-^* mice. **A** Mouse gait was measured by footprint analysis on a straight track, with forepaws in red and hindpaws in blue. **B** There was no significant difference in base width (forepaws, p = 0.775; hindpaws, p = 0.229) or stride length (p = 0.252) during normal walking between *Trim67^-/-^* and *Trim67^+/+^* mice. **C** Latency to fall from an accelerating rotarod in three trials separated by 45 seconds (1, 2, 3), followed by two additional trials 48 hours later separated by 45 seconds (4, 5). Deletion of *Trim67* led to a decrease in fall latency during the first day of trials (p = 0.0074), however there was no difference during the retest trials. **D** Muscle tone measured during a four-paw rolling wire hang assay, reported as maximum and average impulse (weight * time) over 3 trials. *Trim67^-/-^* mice have lower maximum (p = 0.0387) and average (p = 0.0339) hanging impulse than *Trim67^+/+^* mice. * - p < 0.05.

### Deficits in Social Novelty Preference, but not Sociability

Mice were evaluated for the effects of *Trim67* deficiency on social preference using a 3-chamber choice test. In the initial habituation phase, both *Trim67^+/+^* and *Trim67^-/-^* mice made similar numbers of entries into each side chamber (*Trim67^+/+^;* 10.3 ± 1 right, 9.3 ± 0.7 left: *Trim67^-/-^;* 8.5 ± 0.9 right, 8.3 ± 0.8 left). In the test for sociability **(Fig. 7*A***), both genotypes showed a significant preference for spending time in proximity to the cage containing stranger mouse 1 versus an empty cage. However, in the subsequent social novelty phase **(Fig. 7*B***), *Trim67^-/-^* mice showed no preference for a novel stranger mouse, whereas *Trim67^+/+^* mice showed a significant preference. In both sociability and novelty phases, both groups of mice showed a similar number of entries into each side chamber, indicating that the lack of social novelty preference in the *Trim67^-/-^* group was not due to alterations in activity during the test.

**Fig. 7:**
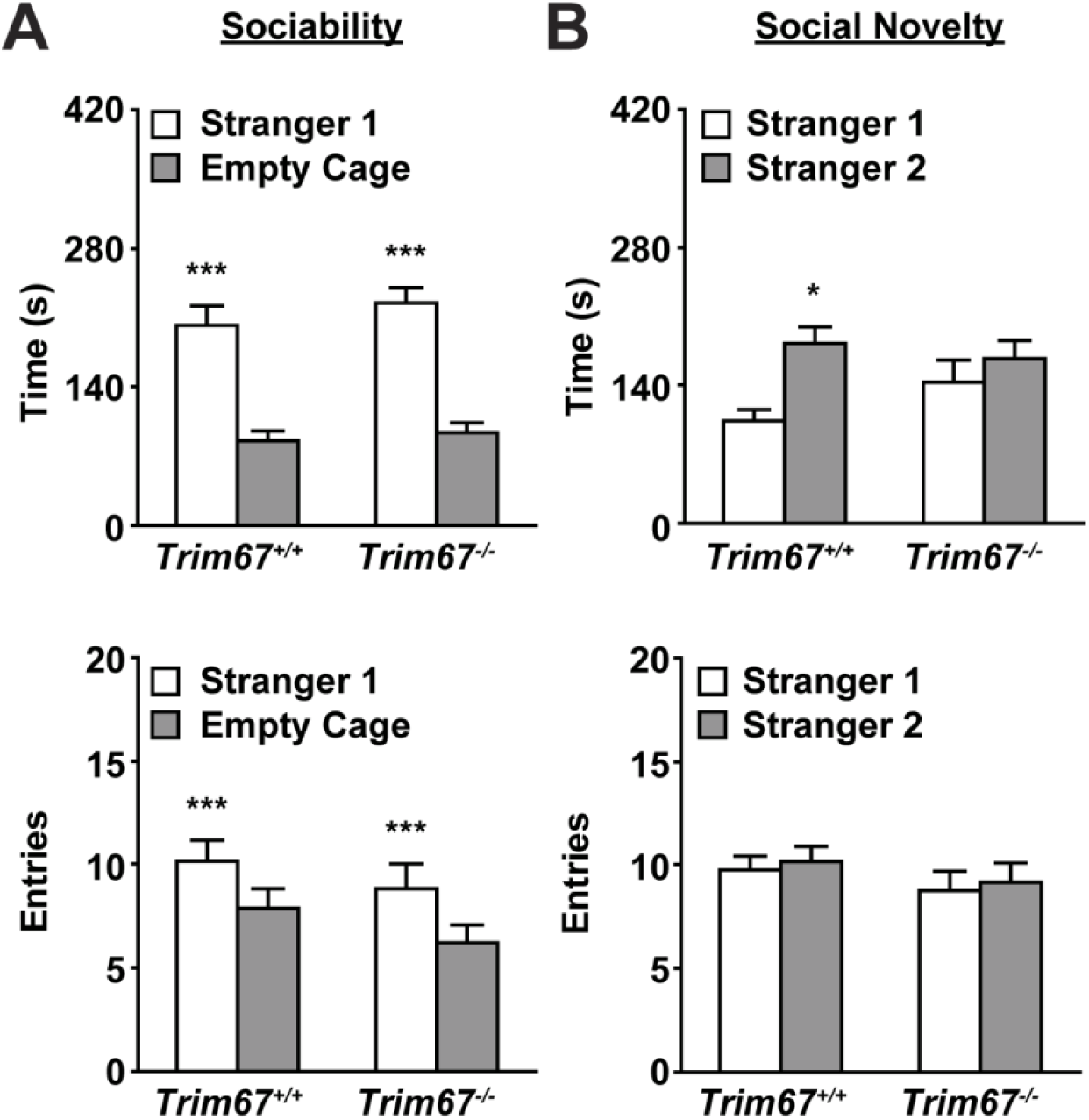
*Trim67^-/-^* mice display normal sociability, but impaired social novelty preference. **A** Measures of total time spent in (top) and entries into (bottom) side chambers of a 3-chamber socialization assay, showing mice of both genotypes exhibited a significant preference for proximity to stranger mouse 1 (*** p < 0.0001). **B** Total time spent in and entries into side chambers in subsequent 3-chamber social novelty assay. *Trim67^-/-^* mice show a lack of preference for the novel stranger, whereas *Trim67^+/+^* mice show a preference (* p = 0.0295).

### Impaired Spatial Memory and Cognitive Flexibility

The Morris water maze was used to assess spatial and reversal learning, swimming ability and visual function. The procedure was divided into three phases: a visible platform test, acquisition with a hidden platform, and reversal learning. In the visible platform trials both groups showed similar times to escape, as well as similar swimming speeds (Table S2). On the first day of hidden platform acquisition both groups again had similar swimming speeds, however *Trim67^-/-^* mice swam significantly more slowly on the first day of the reversal learning test [p = 0.0256].

Following the visible platform task, mice were evaluated for their ability to find a submerged, hidden escape platform. Each animal was given 1 minute per trial to swim to the hidden platform; the criterion for learning was an average group latency to escape of 15 seconds or less **(Fig. 8*A,C*** dashed line). In the acquisition phase, *Trim67^-/-^* mice took significantly longer to find the hidden platform, and never reached the criterion for learning by day 9 of testing **(Fig. 8*A***). On trial day 9, two *Trim67^-/-^* mice failed to locate the platform on more than one trial, and were not tested further. Following the acquisition phase the platform was removed and mice were given a one-minute probe trial. Percent time spent in the quadrant that previously held the platform was significantly higher than the opposite quadrant in *Trim67^+/+^* mice, however *Trim67^-/-^* mice failed to show quadrant selectivity and had lower percent time than *Trim67^+/+^* in the target quadrant [genotype x quadrant interaction, p = 0.0343] **(Fig. 8*B***). *Trim67^-/-^* mice also demonstrated impairment in the number of crossings over the platform’s previous location [main effect of genotype, p = 0.0101, genotype x quadrant interaction, p = 0.0281].

**Fig. 8:**
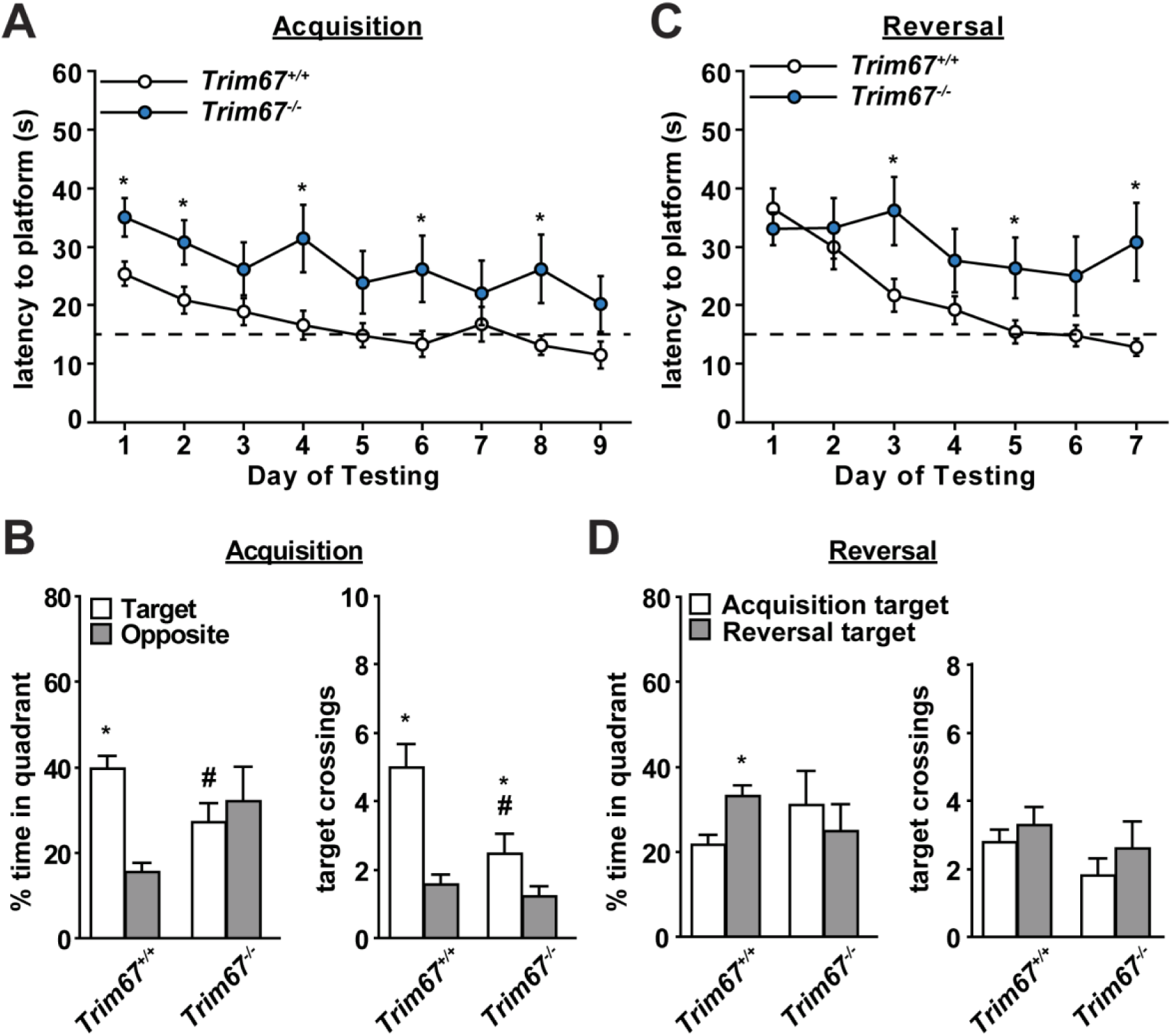
Loss of *Trim67* leads to impairments in spatial learning and memory. **A** Time (s) for mice to find a hidden platform in the Morris water maze, with a threshold for learning set at a group average of 15 s (dashed line). *Trim67^-/-^* mice fail to reach this threshold out to the maximum of 9 days of training, and have overall significantly higher latencies to find the platform (* p<.05, all trials p = 0.0225). **B** Time spent in target and opposite quadrants and number of times crossing the previous platform position in a probe trial following acquisition day 9. Whereas *Trim67^+/+^* mice spent a higher amount of time in the quadrant previously occupied by the platform (* p = 0.0003), *Trim67^-/-^* mice failed to show this preference and spent a lower time in the target quadrant than *Trim67^+/+^* mice (# p = 0.0343). **C** Subsequent reversal trials demonstrated an increased latency to find the platform in the *Trim67^-/-^* cohort compared to *Trim67^+/+^* littermates (* p < 0.05, all trials p = 0.0149). **D** In a probe trial following reversal day 7, only *Trim67^+/+^* mice spent more time in the new target quadrant (* p = 0.0335).

After the acquisition period and probe trials, the platform was placed in a different quadrant and mice were tested until the group reached the learning criterion to assess cognitive flexibility. During this reversal phase of testing, *Trim67^-/-^* mice showed significant deficits in learning the new location of the escape platform [p = 0.0388] **(Fig. 8*C***). Following the last day of reversal testing, the platform was once again removed and mice were given a one-minute probe trial. During this probe trial, only the *Trim67^+/+^* mice showed a preference for spending more time in the quadrant previously containing the platform in the reversal trials, while neither genotype displayed a significant difference in target crossings **(Fig. 8*D***).

### Impairments in Sensorimotor Gating

Mice were tested both at 10–11 weeks and at 17–19 weeks of age for prepulse inhibition of acoustic startle responses. At the first test, *Trim67^-/-^* mice showed no difference in either initial startle amplitude or in the % inhibition by prepulses of any intensity **(Fig. 9*A***). However, in the retest at 17–19 weeks *Trim67^-/-^* mice exhibited a mild decrease in startle response, and significant deficits in prepulse inhibition **(Fig. 9*B***). This suggests impairments in sensorimotor gating emerged by the age of 4 months.

**Fig. 9:**
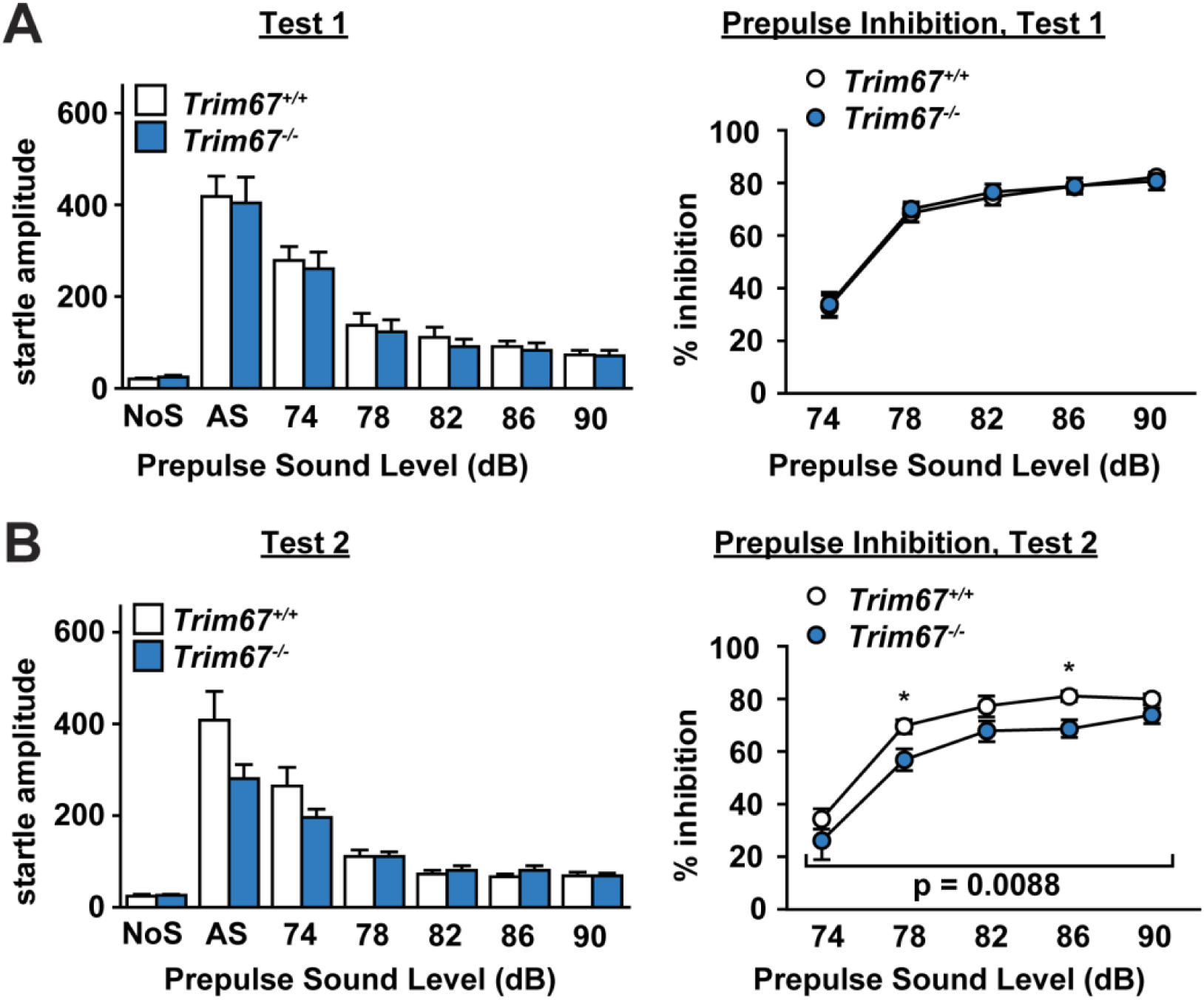
Prepulse inhibition of acoustic startle is impaired by loss of *Trim67*. **A** Startle amplitude (in arbitrary units) in response to no stimulus (NoS) or 120 dB stimulus (AS), with varying prepulse levels 100ms before stimulus. **B** Prepulse inhibition is reported as % inhibition compared to AS. Test 1 took place at 10–11 weeks of age, and Test 2 at 17–19 weeks. *Trim67^-/-^* mice showed a decrease in prepulse inhibition in the second trial (p = 0.0088), but not in the first.

### Altered Subcortical Brain Anatomy in *Trim67^-/-^* Mice

Because TRIM67 is enriched in the brain during development and adulthood, and behavioral testing showed mostly cognitive, memory and social defects, we assessed overall brain anatomy in five-week old mice. Superficially, *Trim67^-/-^* brains appeared smaller than *Trim67^+/+^* counterparts, and were ~10% smaller by post-fixation weight **(Fig. 10*A***). *Trim67^-/-^* brains were matched with *Trim67^+/+^* littermates to reduce variation, sectioned, and stained for myelin. Due to the behavioral phenotypes in *Trim67^-/-^* mice, we measured the size of the hippocampus, lateral ventricle, motor and somatosensory cortex and lateral/basolateral amygdalae **(Fig. 10*B***).

**Fig 10:**
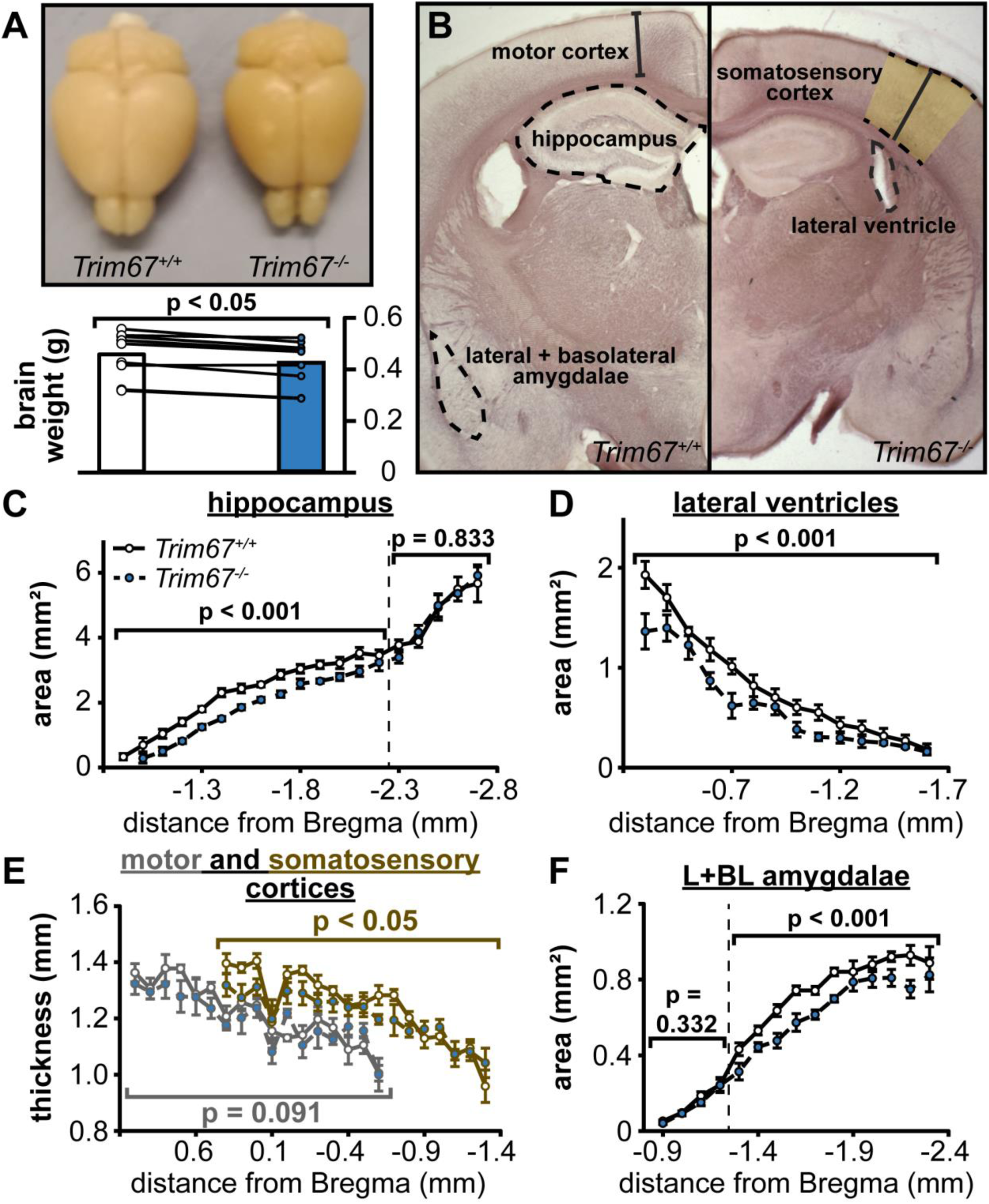
Reduction in total brain weight and size of multiple brain areas occurs with loss of *Trim67*. **A** Photograph of representative *Trim67^+/+^* (left) and *Trim67^-/-^* (right) brains from 5-week old mice. Total weight of *Trim67^-/-^* brains was reduced by approximately 10% when compared to littermate controls (p = 0.0108, Wilcoxon Signed Rank test). **B** Photomicrographs of black-gold stained coronal sections of *Trim67^+/+^* and *Trim67^-/-^* brains at 1.3 mm posterior to Bregma. Outlines delineate regions used for measures in this figure. **C** Area of the hippocampal gray matter (including both the dentate gyrus and Ammon’s horn), reported as the average of both hemispheres from its first emergence through the fimbriae to 2.7 mm posterior to Bregma. The dorsal/anterior hippocampus was smaller in *Trim67^-/-^* brains (p = 0.000110), whereas the first 0.5 mm of the posterior/ventral hippocampus was not different (p = 0.833). **D** Area of the lateral ventricles reported as the average of both hemispheres, from the first section posterior to the anterior commissure to the last section with visible ventricle. *Trim67^-/-^* lateral ventricles were smaller (p = 0.000128). **E** Thickness of the motor and somatosensory cortices, measured from the cingulum to the cortical surface. Deletion of *Trim67* had no effect on the thickness of the motor cortex (p = 0.091), but reduced the thickness of the somatosensory cortex (p = 0.0131). F Area of the combined LA and BLA, reported as the average of both hemispheres. The region before the appearance of the pBLA showed no difference in size with deletion of *Trim67* (p = 0.332), however the remainder of the distinguishable amygdala was smaller (p = 0.000116).

Changes in the size of the hippocampus and lateral ventricles have been associated with spatial learning and memory deficits (Wright et al., 2000; Gaser et al., 2004; Boyer et al., 2007). The anterior portion of the hippocampus was smaller in *Trim67^-/-^* brains **(Fig. 10*C***), whereas the posterior portion was not different between genotypes. Additionally, the anterior aspect of the hippocampal gray matter began approximately 100 μm posterior in *Trim67^-/-^* mice compared to *Trim67^+/+^* mice, when aligned anatomically (Bregma -1.0 and -0.9, respectively). Surprisingly, the lateral ventricles were smaller in *Trim67^-/-^* brains **(Fig. 10*D***), despite the decrease in the size of the hippocampus. The thickness of the somatosensory cortex was also decreased in the brains of *Trim67^-/-^* mice, although to a lesser degree. The motor cortex, however, was the same thickness in both *Trim67^+/+^* and *Trim67^-/-^* brains, suggesting size differences in the hippocampus and ventricles were not due to general hypotrophy **(Fig. 10*E***).

Lesions of the amygdala have been associated with impairments in sensorimotor gating, specifically the basolateral region (Wan and Swerdlow, 1996; Howland et al., 2007). We quantified the combined area of the lateral (LA), anterior basolateral (aBLA), and posterior basolateral amygdalae (pBLA), beginning from the first appearance of the anterior aspect through the external capsule. There was no difference between *Trim67^+/+^* and *Trim67^-/-^* brains in the anterior region before the appearance of the pBLA, however the posterior region was smaller in *Trim67^-/-^* mice **(Fig. 10*F***).

Malformation of the striatum, especially the caudate putamen (CPu) **(Fig. 11*A***), has been associated with deficits in motor and spatial learning (Oliveira et al., 1997; Dang et al., 2006; Lee et al., 2014). The CPu anterior to the globus pallidus was smaller in *Trim67^-/-^* brains, but in the posterior region there was no difference between littermates **(Fig. 11*B***). We generated a binary mask of the myelin-stained CPu at Bregma to determine if the change in CPu was due to a loss of white and/or gray matter **(Fig. 11*C***). Although there was no difference in total area of white matter between genotypes *(Trim67^+/+^* = 2.28 ± 0.12 mm^2^, *Trim67^-/-^* = 2.47 ± 0.12 mm^2^, p = 0.322), when normalized to CPu area, white matter was increased in *Trim67^-/-^* brains *(Trim67^+/+^* = 38.15% ± 1.81%, *Trim67^-/-^* = 46% ± 2.23%, p = 0.030), indicating gray matter was reduced in *Trim67^-/-^* brains. Despite equal white matter in the striatum, the cross-sectional area of the internal capsule, where these fibers fasciculate, was smaller in *Trim67^-/-^* brains **(Fig. 11*D***).

**Fig. 11:**
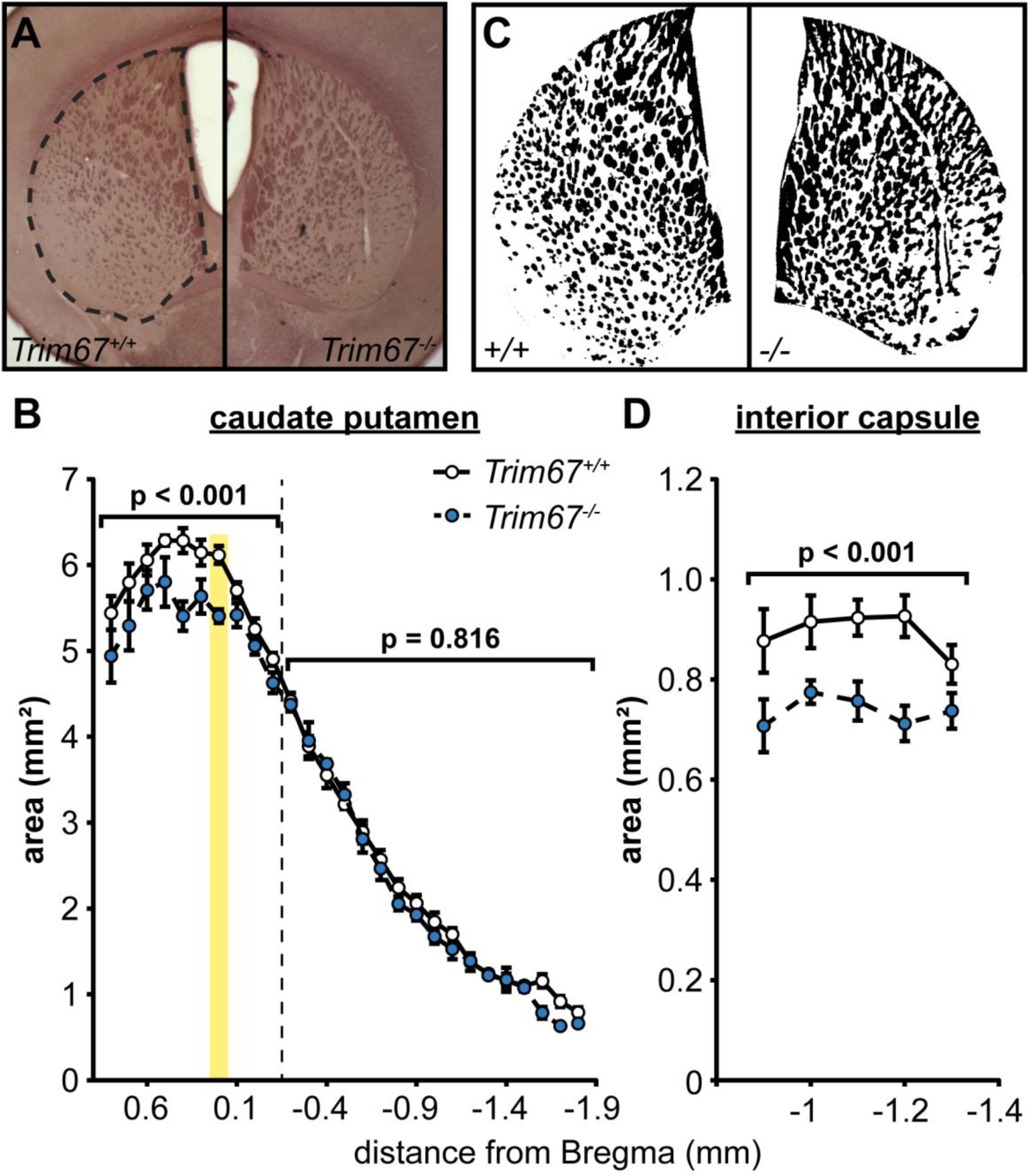
Malformation of caudate putamen in *Trim67^-/-^* mice. **A** Photomicrographs of blackgold stained coronal brain sections showing the area used for quantification of CPu parameters. **B** The area of the CPu, reported as the average of both hemispheres. Yellow box denotes position of white matter measurements. The region of the CPu rostral to the anterior aspect of the globus pallidus was smaller in *Trim67^-/-^* brains (p = 0.000112), however the caudal region was not different (p = 0.816). **C** Binary masks of white matter (black regions) in the CPu after semiautomated segmentation. **D** Deletion of *Trim67* resulted in a decrease in the area of the internal capsule, reported as the average of both hemispheres (p = 0.000138).

Deficits in sensorimotor gating have been associated with abnormalities in the sensory nuclei of the thalamus (Young et al., 1995; Campbell et al., 2007; Hazlett et al., 2008) and TRIM67 expression was evident in this region in the embryonic brain **(Fig. 3*C***). DCC is also present in the fibers of the stria medullaris and habenula (Shu et al., 2000), therefore we measured the size of the thalamus along with some associated fiber tracts **(Fig. 12*A***). The overall size of the thalamus was decreased in the brains of *Trim67^-/-^* mice **(Fig. 12*B***). However, the habenula was the same size compared to *Trim67^+/+^* brains **(Fig. 12*C***), indicating there was not hypotrophy of all diencephalic structures. The stria medullaris and stria terminalis were both smaller in crosssectional area in *Trim67^-/-^* brains **(Fig. 12*D***).

**Fig. 12:**
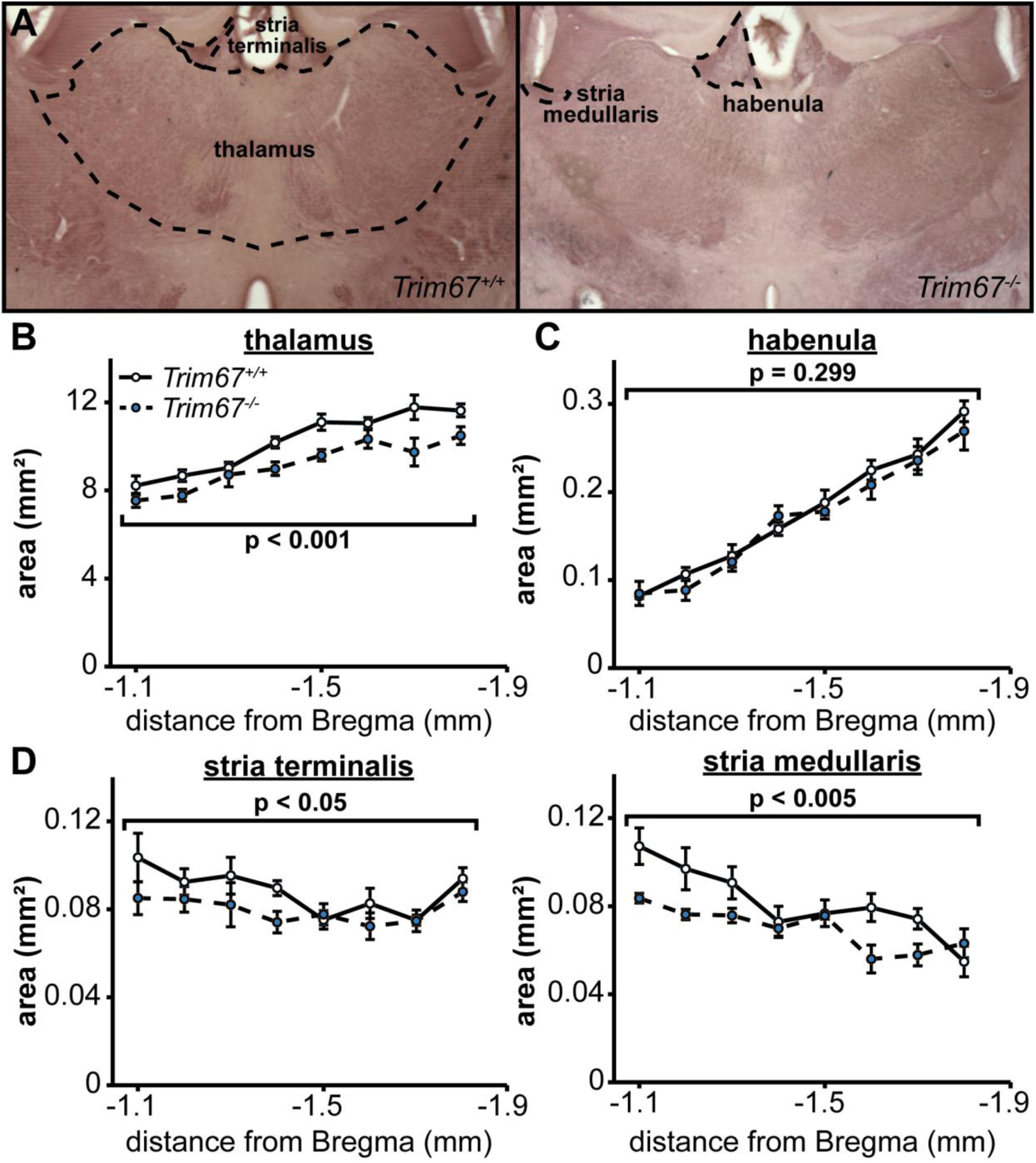
Deletion of *Trim67* is associated with a reduction in thalamus size. **A** Coronal brain sections with the thalamus, habenula, stria terminalis and stria medullaris regions outlined as used for measurements. **B** *Trim67* deletion resulted in a decrease in the area of the thalamus in the region under the anterior hippocampus (p = 0.000174). **C** The area of the habenula was not affected by deletion of *Trim67* (p = 0.299). **D** Both the stria terminalis and stria medullaris were reduced in cross-sectional area in *Trim67^-/-^* brains (p = 0.0222 and 0.00152, respectively).

## DISCUSSION

Here we generated a novel mouse and specific antibody to investigate the largely unstudied E3 ubiquitin ligase, TRIM67. We have shown that TRIM67 is enriched in the cerebellum and, to a lesser extent, other brain regions in the adult mouse. Furthermore, in the embryo TRIM67 is enriched in the developing cortex, diencephalon and midbrain. We found that *Trim67* deletion results in a complex set of behavioral deficits and malformation of several brain regions and axonal fiber tracts in adult mice. In humans, the *TRIM67* gene, which thus far has not been linked to any human disorders, is within the q42.2 band of chromosome 1, a region highly associated with heritable neurological conditions (St. Clair et al., 1990; Millar et al., 2000; Williams et al., 2009). The proximity of *TRIM67* to other genes linked to Alzheimer’s disease and schizophrenia *(DISC1)* (Beecham et al., 2009; Carless et al., 2011), Parkinson’s disease (*SIPA1L2*) (Nalls et al., 2014), and cognitive function *(DISC2, C1orf131, GALNT2)* (Xu et al., 2017) by genome-wide association studies (GWAS), and the behavioral deficits revealed here associated with deletion of murine *Trim67*, may suggest possible involvement of *TRIM67* in human disorders or behaviors. Indeed, a small GWAS of patients with neuroticism, a personality trait that often occurs with major depression and anxiety disorders, identified variations in several genes, including *TRIM67* (Shifman et al., 2008). However, none of these variations reached genome wide significance levels, potentially due to insufficient size of the study.

### A role for TRIM67 in brain development and maintenance

The change in TRIM67 protein levels from embryonic to adult brains suggests that TRIM67 could function both in developing neurons, for example the embryonic cortex, as well as mature neurons, such as Purkinje cells of the adult cerebellum. Indeed, the appearance of sensorimotor gating deficits in prepulse inhibition of acoustic startle at 17–19 weeks, but not at 10–11 weeks, suggests that TRIM67 is necessary for maintenance of brain function in mature animals. The spatial and motor learning impairments also suggest a role for TRIM67 in remodeling of the hippocampus, classically associated with the Morris water maze task (Clark et al., 2008). This study may support an intriguing possibility that the lack of TRIM67 results in progressive neurological dysfunction with age, a common feature of many human disorders including schizophrenia, bipolar disorder and neurodegenerative diseases.

There is apparent hypotrophy of the hippocampus, striatum, thalamus, and parts of the amygdala in *Trim67^-/-^* brains which, based on black-gold staining, is due to decreased non-myelinated tissue. Given the normal size of the motor cortex and some portions of the affected nuclei, this may be due to abnormal maturation and development of specific types of neurons as opposed to overall proliferation. This distinction will require additional studies to determine which neuron types express TRIM67, and at what points during development or maturity TRIM67 is expressed. The pattern of TRIM67 expression in the embryonic brain however supports a role for TRIM67 in neuronal development after proliferation, as TRIM67 protein levels are highest in regions of the developing cortex and hippocampus containing migrating and maturing neurons. This may offer an explanation for the reduction in adult gray matter, as a large portion of this tissue consists of neurites as opposed to cell bodies. A previous study on TRIM67 suggested a role in the formation and elongation of neurites (Yaguchi et al., 2012), and the highly similar protein TRIM9 regulates dendritic arborization in the hippocampus (Winkle et al., 2016), both of which support this possibility.

### A potential role for TRIM67 in netrin-dependent axon development

The decreased gray matter observed in *Trim67^-/-^ mice* alternatively may originate from decreases in branching of axons in the affected brain regions, while the cell bodies of the affected neurons reside in distant nuclei. As netrin can promote axon branching (Dent et al., 2004; Winkle et al., 2014), and TRIM67 interacts with the netrin receptor DCC, this is an intriguing hypothesis. Supporting this possibility, TRIM67 expression is detectable in the nascent substantia nigra embryonically, which projects dopaminergic neurons to the caudate putamen (Matsuda et al., 2009). These axons are guided by netrin-1 expressed in the striatum and branch in response to secreted cues in the same location (Matsuda et al., 2009; Cord et al., 2010; Zhang et al., 2013). The interaction between TRIM67 and both the netrin receptor DCC and TRIM9, an established regulator of axon branching and guidance, suggests a possible role in regulation of netrin-1 dependent axon development. Inhibition of dopaminergic axon branching in the striatum would not only explain the apparent hypotrophy of the caudate putamen, but also deficits in motor function in *Trim67^-/-^* mice. Though less studied, dopaminergic innervation from the substantia nigra and ventral tegmental area is also present in the thalamus (Takada et al., 1990), another region that is smaller in *Trim67^-/-^* brains.

Several netrin-dependent axon tracts are malformed in the brains of *Trim67^-/-^* mice. Though anterior cortical projections are unaffected, the internal capsule and posterior parts of the dorsal commissures are reduced in size. The decrease in internal capsule may seem paradoxical when compared to the comparable amounts of white matter passing through the striatum, however several possibilities may explain this discrepancy. While axons may initially project from the cortex into the striatum, they may not be guided properly to fasciculate fully with the internal capsule. Additionally these axons may not extend to their full length due to a defect in neurite outgrowth rate or persistence, or response to netrin-1, leading to early terminations before reaching the internal capsule. Indeed, such a growth deficit in axons could also contribute to the thinning of the corpus callosum and dorsal hippocampal commissures, and the decreased cross-sectional area of the stria medullaris and stria terminalis. There remains the possibility that these differences are due to overall loss of cells, which may be addressed in future studies.

These experiments have laid a foundation for understanding the role of TRIM67 in development and function of the brain. However, the role of TRIM67 ubiquitin ligase activity, its substrates, and the consequences of their ubiquitination remain to be identified. Further studies will be required to elucidate not only this molecular function of TRIM67 in developing and adult neurons, but also the role of TRIM67 in netrin-dependent responses, the underlying cause of brain hypotrophy resulting from *Trim67* deletion; the time course of and underlying cause of behavioral abnormalities in *Trim67^-/-^* mice; and the contribution of *TRIM67* variations to human neurological disorders. Furthermore, although TRIM67 is the most evolutionarily conserved vertebrate class I TRIM, unlike the invertebrate TRIM ortholog, loss of *Trim67* does not fully phenocopy the axon branching and guidance defects seen in mice lacking *Ntn1* or *Dcc*. Future studies must examine whether other class I TRIMs, such as TRIM9, which interacts with DCC and TRIM67, or TRIM1 and TRIM18, whose mutation in humans is associated with midline birth defects, have coordinated or parallel functions to TRIM67 downstream of netrin/DCC.

## Acknowledgements

We thank Enaj Furigay and Carey Hanlin for experimental assistance, Saumil Patel for assistance with the mouse colony, and Viktoriya Nikolova and Dr. Natallia Riddick for assistance with behavioral assays. This work was supported by National Institutes of Health GM108970 (S.L.G.) and F31NS096823 (N.P.B.).

